# Guardian of Excitability: Multifaceted Role of Galanin in Whole Brain Excitability

**DOI:** 10.1101/2024.04.05.588285

**Authors:** Nicolas N. Rieser, Milena Ronchetti, Adriana L. Hotz, Stephan C. F. Neuhauss

## Abstract

Galanin is a neuropeptide, which is critically involved in homeostatic processes like controlling arousal, sleep, and regulation of stress. This extensive range of functions aligns with implications of galanin in diverse pathologies, including anxiety disorders, depression, and epilepsy. Here we investigated the regulatory function of galanin on whole-brain activity in larval zebrafish using wide-field Ca^2+^ imaging. Combining this with genetic perturbations of galanin signaling and pharmacologically increasing neuronal activity, we are able to probe actions of galanin across the entire brain. Our findings demonstrate that under unperturbed conditions and during epileptic seizures, galanin exerts a sedative influence on the brain, primarily through the galanin receptor 1a (*galr1a*). However, exposure to acute stressors like pentylenetetrazole (PTZ) compromises galanin’s sedative effects, leading to overactivation of the brain and increased seizure occurrence. Interestingly, galanin’s impact on seizures appears to be bidirectional, as it can both decrease seizure severity and increase seizure occurrence, potentially through different galanin receptor subtypes. This nuanced interplay between galanin and various physiological processes underscores its significance in modulating stress-related pathways and suggests its potential implications for neurological disorders such as epilepsy. Taken together, our data sheds light on a multifaceted role of galanin, where galanin regulates whole-brain activity but also shapes acute responses to stress.

## Introduction

Similar to classic neurotransmitters, neuropeptides are chemical messengers that are mainly synthesized and released by neurons. In contrast to conventional neurotransmitters, neuropeptides are amino acid chains between 3 and 36 amino acids of length. While classic neurotransmitters are primarily stored in synaptic vesicles, neuropeptides are predominantly stored in large dense-core vesicles (LDCV) and are mostly released during neuronal bursts or high-frequency firing ^1,2^. Neuropeptides play a crucial role in modulating neuronal activity and regulating various aspects of neural network function. They act as neuromodulators, exerting long-lasting effects on neuronal excitability, synaptic transmission, and plasticity ^1,3^. By influencing the balance between excitatory and inhibitory inputs to neurons, neuropeptides can shape network dynamics and information processing. This is achieved through their interactions with specific receptors and signaling pathways, allowing for precise modulation and optimization of neural activity in response to changing conditions ^2^. Overall, neuropeptides play a pivotal role in orchestrating the complex activity of neuronal networks and maintaining homeostasis within the central nervous system. Elevated neuronal activity is often accompanied by shifts in neuropeptide transcription rates through a phenomenon known as “stimulus-secretion-synthesis coupling” ^4,5^. This intricate interplay between neuropeptide release and biosynthesis likely serves as a crucial mechanism for replenishing neuropeptide stores, given the absence of a mechanism for neuropeptide uptake post-release and their degradation by extracellular peptidases. Further, neuropeptides are well known to contact neurons via volume transmission, where extrasynaptically secreted molecules activate receptors on neurons synaptically unconnected to the releasing neuron. This leads to signal transmission across considerable distances, extending up to multiple microns, targeting multiple neurons in different regions of the brain ^1,2,6–9^.

Galanin is such a neuropeptide, which is expressed predominantly in the hypothalamus, the center of homeostatic regulation in the brain. Galanin is highly involved in homeostatic processes like controlling arousal and sleep ^10–16^, but has been shown to regulate stress ^17–21^. Moreover, research has demonstrated that galanin-producing neurons in the hypothalamus play a role in regulating both food intake ^22–26^ and parental behavior ^27,28^ in rodents. This extensive range of functions aligns with implications of galanin in diverse pathologies, including anxiety disorders, depression, and epilepsy ^1,18,19,29–35^. We found an upregulation of galanin in a recently described novel model of epilepsy^36^ that let us hypothesize that galanin may mediate a neuroprotective, net inhibitory effect on epileptic brains.

Here we investigated the regulatory function of galanin on whole-brain activity in larval zebrafish (*Danio rerio*). Leveraging the transparency of zebrafish during their larval stages, a characteristic that facilitates live imaging, we utilized basic wide-field Ca^2+^ imaging methods. Combining this with genetic perturbation of galanin and pharmacologically increasing neuronal activity, we were able to delve into the actions of galanin across the entire brain.

Our findings demonstrate that, under unperturbed conditions and during epileptic seizures, galanin exerts a net inhibitory influence on the brain, which is likely governed by *galanin receptor 1a* (*galr1a*). However, when faced with an acute stressor like pentylenetetrazole (PTZ), the typically sedative effects of galanin are substantially compromised. We found that exposure to this acute central nervous system stressor results in a galanin dependent overactivation of the brain that overrides most of galanins sedating actions and increases the occurrence of epileptic seizures. Taken together, our data sheds light on a multifaceted role of galanin, where galanin regulates whole-brain activity but also shapes acute responses to stress.

## Results

### *gal* expression correlates with whole-brain activity

In prior work, we introduced a novel epilepsy model characterized by recurrent epileptic seizures and interictal neuronal hypoactivity (Figure 1A - D) ^36^. The unexpected observation of locomotor and neuronal hypoactivity in this model stands in contrast to most existing studies in zebrafish that report hyperactivity in epileptic animals ^37–39^. Intriguingly, recent investigations have demonstrated that overexpression of galanin induces locomotory hypoactivity in zebrafish models ^10,11^. To explore the potential involvement of galanin in the hypoactivity observed in *eaat2a* mutants, we conducted qPCR analysis on 5 dpf (days post fertilization) larval brains (Figure 1C). The results revealed a significant 15.4-fold increase in galanin expression compared to wild type siblings, suggesting an involvement of galanin in th interictal hypoactivity observed in *eaat2a* mutants. Furthermore, recent findings have shed light on the role of galanin in pharmacologically induced rebound sleep ^12^. Additionally, it was demonstrated that increasing short term neuronal activity results in subsequent inactivity that is dependent on galanin ^12^. To explore whether the application of GABA^A^ receptor antagonist pentylenetetrazole (PTZ) would also lead to a temporary decrease in whole-brain activity, we exposed 5 dpf larvae to 20 mM PTZ for 1 h, followed by a 2-hour washout period before undergoing Ca^2+^ imaging. Remarkably, while PTZ increases swimming activity and induces seizure-like behavior during acute drug exposure ^12,39,40^, there was a significant reduction in brain activity during PTZ rebound (Figure 1G – K), which was correlated with an increase in galanin expression by 2.5-fold (Figure 1I) compared with non-PTZ-treated larvae. Notably, large Ca^2+^ fluctuations (ΔF/F^0^ > 10%) decreased in frequency (Figure 1J), and decreased in amplitude (Figure 1K), while their duration increased (Figure 1L) compared to non-PTZ-treated larvae. For small Ca^2+^ fluctuations (ΔF/F^0^ > 5) the frequency also decreased (Figure 1J), while their amplitude and duration were not affected (Figure 1K and L). Together, our results illustrate that the expression of galanin rises in reaction to seizure-like activity in two separate seizure models and is correlated with an overall decrease in whole-brain activity.

**Figure 1.**
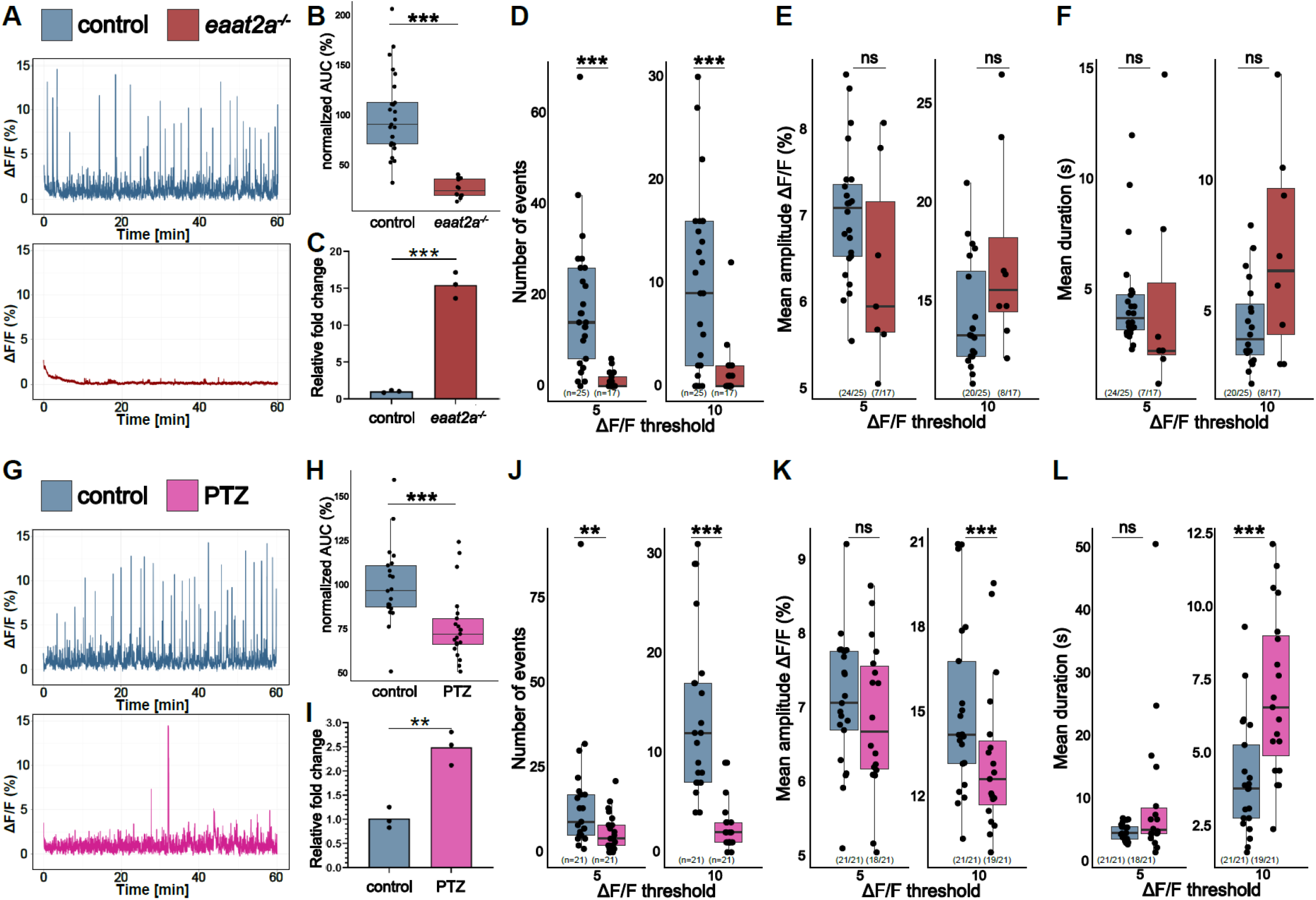
*gal* Expression Correlates with Whole-Brain Activity. A) Representative calcium signals (*elavl3:GCaMP5G*) recorded across the brain of 5 dpf control (*eaat2a*^*+/+*^) larva (blue, top) and *eaat2a*^*-/-*^ mutant without seizure activity (red, bottom). B) AUC calculated and averaged over two five-minutes time windows per animal and normalized to control (n control = 25; n *eaat2a*^*-/-*^ = 12). C) Galanin transcript levels in pools of control vs. *eaat2a*^*-/-*^ brains (n = 3). D) Number of Ca^2+^ events above 5% ΔF/F_0_ (left) and 10% ΔF/F_0_ (right). E) Average amplitude of Ca^2+^ events above 5% ΔF/F_0_ (left) and 10% ΔF/F_0_ (right) per larva. F) Average duration of Ca^2+^ events above 5% ΔF/F_0_ (left) and 10% ΔF/F_0_ (right) per larva. G) Representative calcium signals (*elavl3:GCaMP6f*) recorded across the brain of 5 dpf control (blue, top) and larvae 2 hours after PTZ exposure (PTZ rebound) (magenta, bottom). H) AUC calculated over 1 h recording, normalized to control (n control = 21; n PTZ rebound = 21). I) Galanin transcript levels in pools of control vs. PTZ rebound brains (n = 3). G) Number of Ca^2+^ events above 5% ΔF/F_0_ (left) and 10% ΔF/F_0_ (right). J) Average amplitude of Ca^2+^ events above 5% ΔF/F_0_ (left) and 10% ΔF/F_0_ (right) per larva. K) Average duration of Ca^2+^ events above 5% ΔF/F_0_ (left) and 10% ΔF/F_0_ (right) per larva. Significance levels: ***p < .001, **p < .01, *p < .05, ns = not significant (p > .05), Wilcoxon-Mann-Whitney test (B, E, H, J, K), negative binomial regression (D, I), Student’s t-test (C, F).

### *gal* controls whole-brain activity

To investigate if galanin by itself is able to reduce whole-brain activity, we employed a transgenic approach by using *hsp70l:gal* ^10^ expressing larvae. These transgenic fish express galanin under the control of the heatshock promoter *hsp70l*, enabling the induction of galanin expression by upregulating transcripts by over 300 fold ^11^. Attempts to induce galanin overexpression through heat shock were discontinued due to a notable impact on the brain activity of wild type larvae compared to non-exposed wild type larvae (Suppl. Figure 1). Yet, considering that *hsp70l:gal* larvae already display a basal elevation of galanin transcripts by approximately 8-fold ^11^, comparable with the upregulation seen in *eaat2a*^*-/-*^ mutants and PTZ rebound (Figure 1C and 1I), we chose to use larvae without inducing transcription via heat shock. We performed Ca^2+^ imaging and found that indeed elevated expression of galanin in *hsp70l:gal* larvae leads to a decrease in whole-brain activity (Figure 2A) compared to wild type siblings. We detected a reduction in frequency of Ca^2+^ fluctuations for larger Ca^2+^ events and a reduction in their amplitude (Figure 2D and 2E), mirroring our earlier findings in the PTZ rebound (Figure 1G and 1H). Notably, *hsp70l:gal* larvae exhibited no difference in small Ca^2+^ fluctuations (Figure 2D – 2F). Subsequent quantification of galanin levels in the brain through qPCR unveiled a 1.8-fold increase in galanin transcripts (Figure 2C) in *hsp70l:gal* larvae compared to their wild type siblings, potentially explaining the more subtle effects observed on brain activity. To further examine the role of galanin in regulating whole brain activity, we used *galanin*^*t12ae/t12ae*^ mutants (*gal*^-/-^ mutants) that carry a 7 bp deletion in exon 3 of the *galanin* gene which leads to a premature stop codon 41. The absence of galanin protein was confirmed in an antibody staining showing a characteristic half-ring of galanin-positive fibers in the diencephalon that was completely absent in *gal*^-/-^ mutants (Suppl. Figure 2). While other studies focused their analysis in *gal*^-/-^ mutants on their locomotion ^12,17^ and sleep ^12^, we want to assess the effects of a galanin deficiency on the brain. When performing Ca^2+^ imaging, we found that brain activity is heavily increased in *gal*^*-/-*^ mutants (Figure 2G) compared to wild type larvae. While overexpression of galanin primarily influences larger Ca^2+^ events, the absence of galanin specifically affects smaller Ca^2+^ fluctuations. The increased frequency of these events is accompanied by a decrease in both amplitude and duration (Figure 2I – 2K). To delve deeper into the galanin system, we took a targeted approach focusing on a specific galanin receptor in zebrafish. Zebrafish have four galanin receptors: *galr1a, galr1b, galr2a*, and *galr2b*. While information about these receptors in zebrafish is limited, studies indicate that *galr1a* is widely expressed in the brain, and sequence identity closely resemble those of mammalian GALR1 ^41–43^. To investigate the actions of *galr1a* in zebrafish, we conducted F0 knockouts which enable a direct analysis of the injected embryos, bypassing the need to establish stable lines ^44^. We found that *galr1a* crispants have an increased number of small Ca^2+^ events in the brain, compared to control injected larvae (Figure 2O). The absence of *galr1a* specifically affects the frequency of smaller Ca^2+^ fluctuations, similar to *gal*^*-/-*^ mutants (Figure 2I), suggesting that galanin might regulate whole-brain activity via *galr1a*. We could not detect an upregulation of galanin in *galr1a* crispants, ruling out genetic compensation effects (Figure 2N).

Taken together, we could show that galanin is crucial in controlling spontaneous brain activity, and that this is at least partially regulated by *galr1a*.

**Figure 2.**
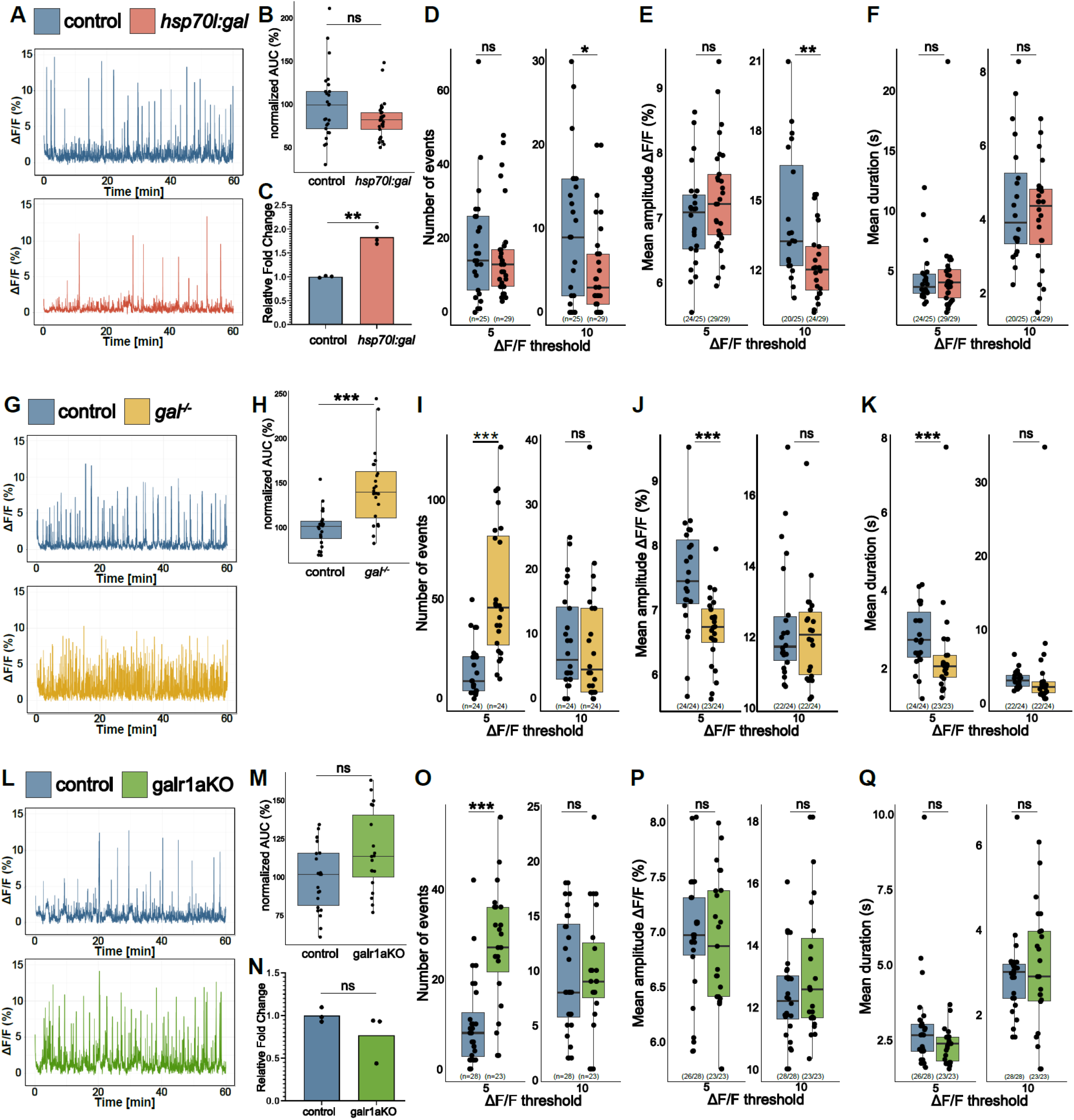
*gal* Controls Whole-Brain Activity. A) Representative calcium signals (*elavl3:GCaMP5G*) recorded across the brain of 5 dpf control (blue, top) and *hsp70l:gal* sibling (orange, bottom). B) AUC calculated over 1 h recording, normalized to control (n control = 25; n *hsp70l:gal* = 29). C) Galanin transcript levels in pools of control vs. *hsp70l:gal* brains (n = 3). D) Number of Ca^2+^ events above 5% ΔF/F_0_ (left) and 10% ΔF/F_0_ (right). E) Average amplitude of Ca^2+^ events above 5% ΔF/F_0_ (left) and 10% ΔF/F_0_ (right) per larva. F) Average duration of Ca^2+^ events above 5% ΔF/F_0_ (left) and 10% ΔF/F_0_ (right) per larva. G) Representative calcium signals (*elavl3:GCaMP5G*) recorded across the brain of 5 dpf control (blue, top) and *gal*^*-/-*^ larva (yellow, bottom). H) AUC calculated over 1 h recording, normalized to control (n control = 24; n *gal*^*-/-*^ = 24). I) Number of Ca^2+^ events above 5% ΔF/F_0_ (left) and 10% ΔF/F_0_ (right). J) Average amplitude of Ca^2+^ events above 5% ΔF/F_0_ (left) and 10% ΔF/F_0_ (right) per larva. K) Average duration of Ca^2+^ events above 5% ΔF/F_0_ (left) and 10% ΔF/F_0_ (right) per larva. L) Representative calcium signals (*elavl3:GCaMP6f*) recorded across the brain of 5 dpf control injected (blue, top) and *galr1a* crispants (galr1aKO) larva (green, bottom). M) AUC calculated over 1 h recording, normalized to control (n control = 28; n *galr1a* crispants = 23). N) Galanin transcript levels in pools of control vs. galr1aKO brains (n = 3). O) Number of Ca^2+^ events above 5% ΔF/F_0_ (left) and 10% ΔF/F_0_ (right). P) Average amplitude of Ca^2+^ events above 5% ΔF/F_0_ (left) and 10% ΔF/F_0_ (right) per larva. Q) Average duration of Ca^2+^ events above 5% ΔF/F_0_ (left) and 10% ΔF/F_0_ (right) per larva. Significance levels: ***p < .001, **p < .01, *p < .05, ns = not significant (p > .05), Wilcoxon-Mann-Whitney test (B, E, F, H, J, K, M, O, P), negative binomial regression (D, I, N), Student’s t-test (C).

**Figure 3.**
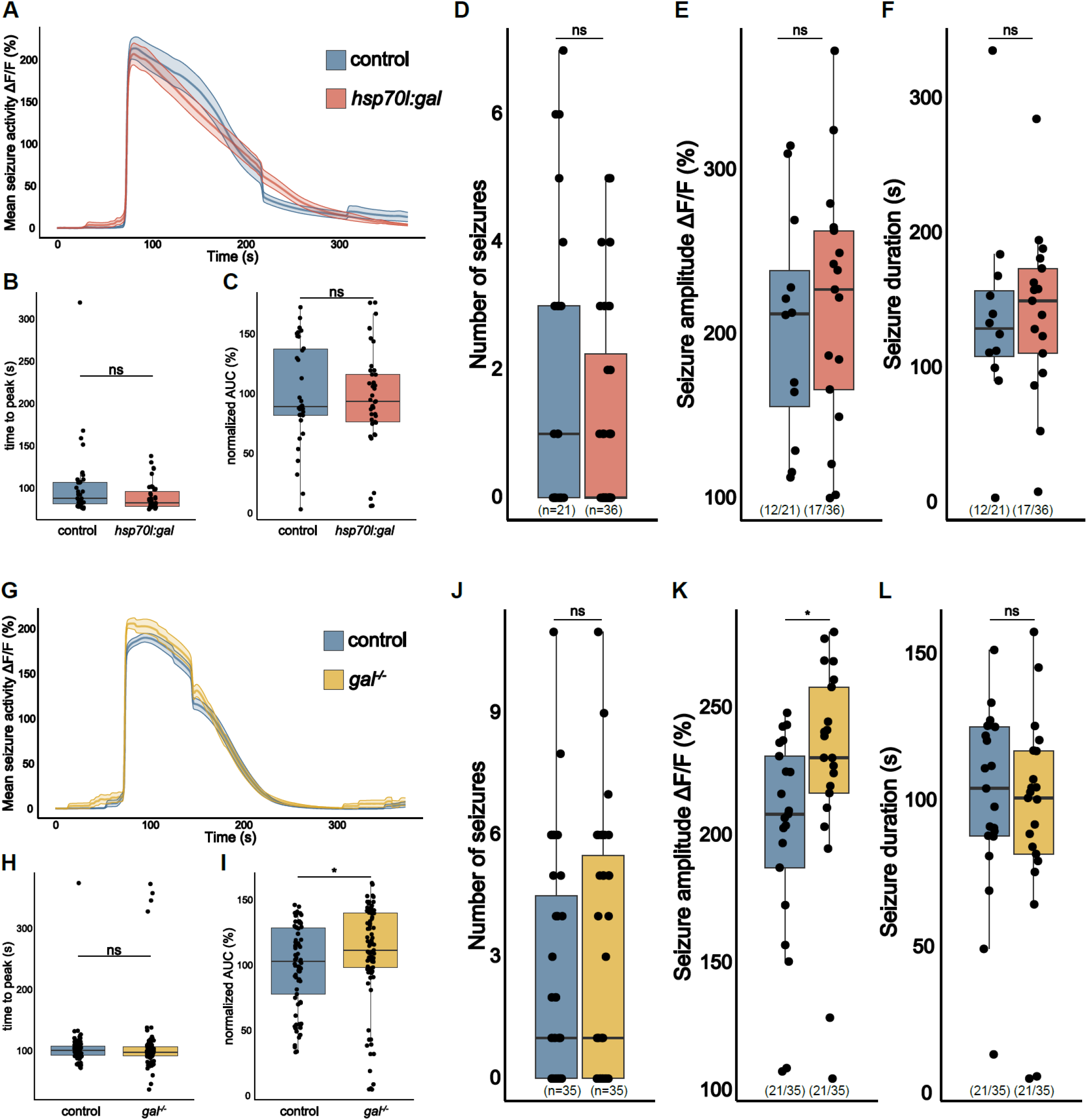
*gal* Exerts a Modest Effect on Seizure Activity in *eaat2a*^-/-^ Mutants. A) Averaged calcium signals (*elavl3:GCaMP5G*) for spontaneous seizures recorded across the brain of 5 dpf control (*eaat2a*^*-/-*^) larva (blue, 35 events) and *eaat2a*^*-/-*^*;hsp70l:gal* (orange, 39 events) aligned by 50% of maximum amplitude. Shaded area represents SEM. B)Time to peak calculated from beginning of aligned seizure until maximum ΔF/F_0_ signal. C) AUC calculated over spontaneous seizures normalized to control (n control = 35; n *eaat2a*^*-/-*^*;hsp70l:gal* = 39). D) Number of spontaneous seizures per larva. E) Amplitude of spontaneous seizures per larva. F) Duration of spontaneous seizures per larva. G) Averaged calcium signals (*elavl3:GCaMP5G*) for spontaneous seizures recorded across the brain of 5 dpf control (*eaat2a*^*-/-*^) larva (blue, 77 events) and *eaat2a*^*-/-*^*;gal*^*-/-*^ (yellow, 89 events) aligned by 50% of maximum amplitude. Shaded area represents SEM. H) Time to peak calculated from beginning of aligned seizure until maximum ΔF/F_0_ signal. I) AUC calculated over spontaneous seizures normalized to control (n control = 77; n *eaat2a*^*-/ -*^ *;gal*^*-/-*^ = 89). J) Number of spontaneous seizures per larva. K) Amplitude of spontaneous seizures per larva. L) Duration of spontaneous seizures per larva. Significance levels: ***p < .001, **p < .01, *p < .05, ns = not significant (p > .05), Wilcoxon-Mann-Whitney test (B, C, E, F, H, I, K, L), negative binomial regression (D, J).

### *gal* exerts only a modest effect on seizure activity in *eaat2a* mutants

While we could show that galanin has direct implications in regulating whole brain activity, others have investigated its antiepileptic properties. It was shown that galanin signaling can affect the seizure thresholds and severity in mice ^32–35^, and can reduce the occurrence of seizure-like behavioral episodes and their intensity in zebrafish ^11^. The increased levels of galanin observed in *eaat2a*^*-/-*^ mutants could be interpreted as an internal compensatory mechanism, possibly serving to shield neurons from excitotoxicity. The observed interictal hypoactivity may be a consequence of the brain’s deliberate neuroprotective strategy so reduce the occurrence of seizures. To investigate if galanin could indeed act as a neuroprotective agent we examined if additional galanin leads to seizure protection in *eaat2a*^*-/-*^ mutants. *eaat2a*^*-/-*^ mutants exhibit global spontaneous seizures lasting all the way up to over 6 minutes in duration ^36^ (Figure 3). Although not all *eaat2a*^*-/-*^ mutants exhibit spontaneous epileptic seizures in the 1-hour recording window, an estimated 57.14% - 60.00% (12/21; 21/35) of larvae do (Figure 3). We then compared seizures of 5 dpf *eaat2a*^*-/-*^ *;hsp70l:gal* with *eaat2a*^*-/-*^ larvae and found no significant difference in seizure susceptibility or severity in spontaneous epileptic seizures. Neither the number of seizures, their amplitude, duration nor time to peak was altered (Figure 3A – 3F). This is however not surprising, as we showed earlier that 5 dpf *eaat2a*^*-/-*^ mutants already overexpress galanin by 15.4-fold (Figure 1C) and therefore more galanin might not make a big difference. To investigate if galanin is still important in seizure protection, we turned to analyze *eaat2a*^*-/-*^*;gal*^*-/-*^ mutants (Figure 3G – L). We found that area under the curve (AUC) of seizures is slightly increased (Figure 3I) and seizure amplitudes are increased compared to *eaat2a*^*-/-*^ controls (Figure 3K). Neither the number of seizures, the seizure duration nor the time to peak changed significantly (Figure 3H, J and L). Taken together, galanin demonstrates only a modest level of seizure protection in *eaat2a*^*-/-*^ mutants.

### *gal* promotes seizures in PTZ exposed larvae

To investigate if galanin could mediate epileptic seizures in another state-of-the-art seizure model, we turned again to the GABAA receptor antagonist, PTZ. We applied 20 mM PTZ to the larvae directly before Ca2+ imaging and compared PTZ exposed larvae with their unexposed siblings. PTZ exposure increases neuronal activity in a progressive manner and triggers short epileptic seizures lasting up to 2 minutes in duration 36 (Figure 4).

Despite the observed decrease in neuronal activity resulting from galanin overexpression (Figure 2A, 2D and 2E), exposing 5 dpf *hsp70l:gal* larvae to 20 mM PTZ surprisingly led to a significant increase in seizure activity (Figure 4D) compared to non-exposed wild type siblings. While seizure number increases significantly (Figure 4D), there is also a decrease in seizure duration (Figure 4F). Conversely, the absence of galanin during acute PTZ exposure induces an opposing phenotype (Figure 4J – K). A marked decrease in the occurrence of epileptic seizures is evident in *gal*^*-/-*^ mutants (Figure 4K) compared to wild type larvae. Merely 6 out of 38 *gal*^*-/-*^ mutants (15.8%) display seizures, whereas 20 out of 39 wild type larvae (51.3%) exhibit seizure activity. However, when *gal*^*-/-*^ larvae experience seizures upon PTZ, the seizure amplitude is significantly reduced (Figure 4K), while their duration (Figure 4L) increases. Further, time to peak and AUC of seizures are increased (Figure 4H – 4I). Strikingly, while seizures in *eaat2a*^*-/-*^ control larvae come to an abrupt stop after a gradual decline, seizures seem to taper off over the duration of several minutes in *gal*^*-/-*^ mutants (Figure 4G).

Contrary to our initial expectations derived from prior results indicating a potentially sedative and neuroprotective role of galanin, the exposure of larvae to PTZ under acute conditions resulted in an opposing observation. While it was surprising that galanin seems to increase seizure susceptibility instead of protecting the larvae from seizures, this fits very well with results from a recent publication, where the authors found galanin to be involved in a stress response upon salient stimuli ^17^. The authors hypothesize that galanin binds to autoreceptors on Gal^+^ neurons, reducing their own activity and therefore reducing their secretion of galanin and additionally reduce release of GABA on downstream stress-promoting neurons, ultimately leading to an increase in stress ^17^. While the specific galanin receptor governing the stress pathway remains elusive, it has been found that GALR1 exerts an inhibitory influence on neuronal activity in mammals, while GALR2 displays a dual role, exhibiting both inhibitory and excitatory effects ^1,45^. Notably, it has been demonstrated that a significant portion of Gal+ neurons in the preoptic area in zebrafish express either *galr1a, galr1b* or *galr2b* ^17^. To discern whether *galr1a*, beyond its role in regulating whole-brain activity, is intricately linked to the stress response pathway, we implemented F0 knockouts of *galr1a* and exposed injected larvae to PTZ-induced stress. If *galr1a* were the autoreceptor on GABAergic Gal+ neurons, we would expect a decrease in seizure number similar to *gal*^*-/-*^ mutants. However, our examination of *galr1a* crispants exposed to 20 mM PTZ revealed no significant decrease in seizure frequency compared to control injected larvae (Figure 4P). This suggests that *galr1a* is not the hypothesized autoreceptor governing this stress response pathway. Furthermore, our findings reveal that *galr1a* crispants manifest a more severe response to PTZ-induced seizures, evident in the substantial increase in both amplitude and duration compared to control injected larvae (Figure 4Q and 4R). Strikingly, several seizure characteristics in *galr1a* crispants mirror those observed in *gal*^-/-^ mutants exposed to PTZ. Specifically, AUC of seizures shows a marked increase, the time to peak extends, and seizures exhibit a gradual tapering off over several minutes, a distinct pattern in contrast to the rapid resolution seen in controls (Figure 4M – 4O).

**Figure 4.**
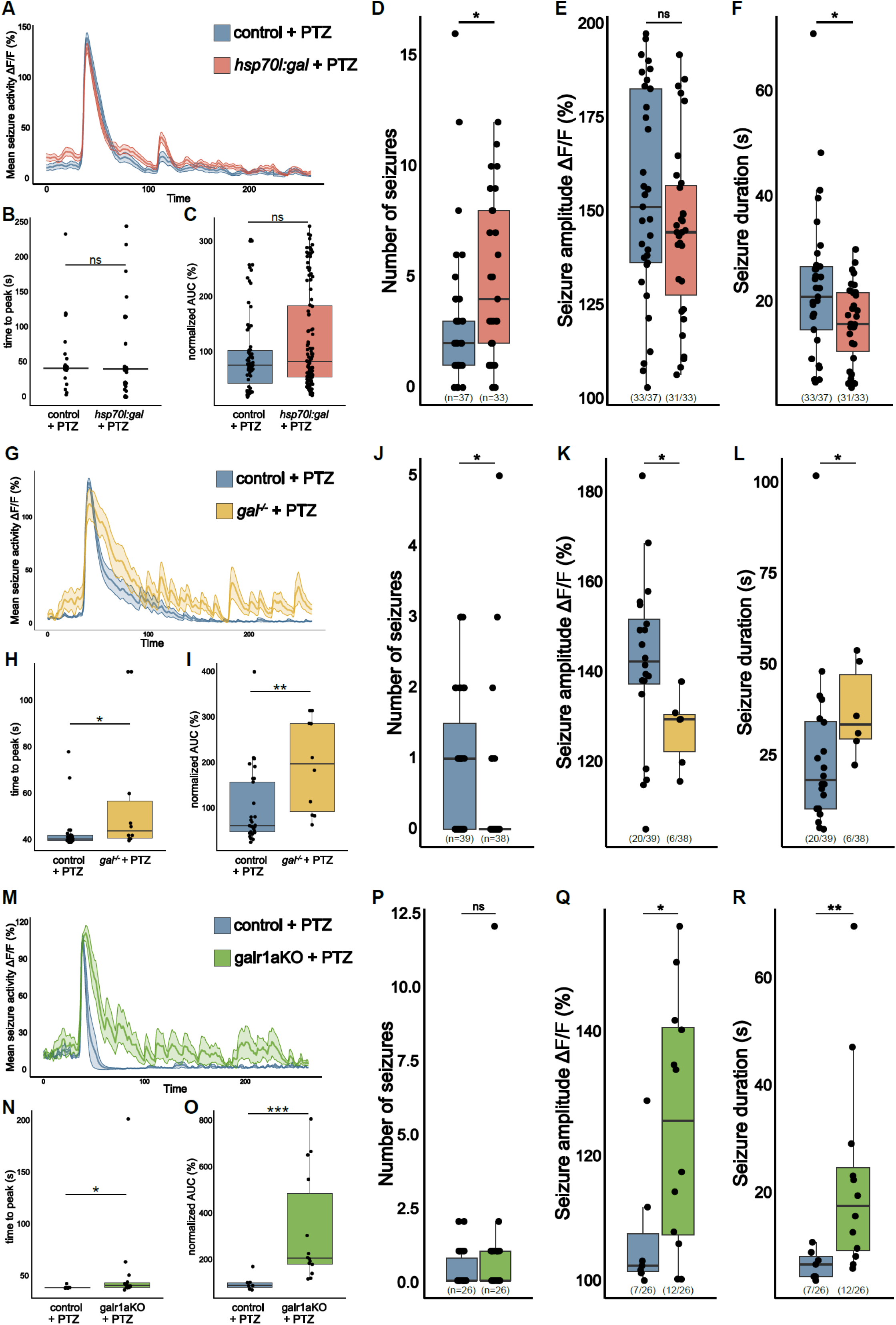
gal Promotes Seizures in PTZ Exposed Larvae. A) Averaged calcium signals (*elavl3:GCaMP6f*) for seizures elicited by 20 mM PTZ recorded across the brain of 5 dpf control (blue, 76 events) and *hsp70l:gal* (orange, 113 events) aligned by 50% of maximum amplitude. Shaded area represents SEM. B) Time to peak calculated from beginning of aligned seizure until maximum ΔF/F_0_ signal. C) AUC calculated over seizures normalized to control (n control = 76; n *hsp70l:gal* = 113). D) Number of seizures per larva. E) Amplitude of seizures per larva. F) Duration of seizures per larva. G) Averaged calcium signals (*elavl3:GCaMP6f*) for seizures elicited by 20 mM PTZ recorded across the brain of 5 dpf control (blue, 29 events) and *gal*^*-/-*^ (yellow, 10 events) aligned by 50% of maximum amplitude. Shaded area represents SEM. H) Time to peak calculated from beginning of aligned seizure until maximum ΔF/F_0_ signal. I) AUC calculated over seizures normalized to control (n control = 29; n *gal*^*-/-*^ = 10). J) Number of seizures per larva. K) Amplitude of seizures per larva. L) Duration of seizures per larva. M) Averaged calcium signals (*elavl3:GCaMP6f*) for seizures elicited by 20 mM PTZ recorded across the brain of 5 dpf control injected (blue, 7 events) and *galr1a* crispants (galr1aKO, green, 14 events) aligned by 50% of maximum amplitude. Shaded area represents SEM. N) Time to peak calculated from beginning of aligned seizure until maximum ΔF/F_0_ signal. O) AUC calculated over seizures normalized to control (n control = 7; n *galr1a* crispants = 14). P) Number of seizures per larva. Q) Amplitude of seizures per larva. R) Duration of seizures per larva. Significance levels: ***p < .001, **p < .01, *p < .05, ns = not significant (p > .05), Wilcoxon-MannWhitney test (B, C, E, F, H, I, K, L, N, O, Q, R), negative binomial regression (D, J, P).

Taken together, our findings underscore the multifaceted role of galanin, not only in inducing a sedating effect on the entire brain but also in playing a pivotal role in a recently described stress response pathway ^17^. Although the specific receptor orchestrating this stress pathway remains unidentified, our research decisively eliminates *galr1a* as the regulator. Collectively, this leads us to the conclusion that galanin likely exerts its sedative actions during epileptic seizures through *galr1a*, and that the stress response pathway is governed by another galanin receptor.

## Discussion

Sustaining brain homeostasis enables consistent behavior and demands the brain’s ongoing integration of sensory information from the environment, internal behavioral state of the organism, and the elaborate networks of synaptic and non-synaptic connectivity ^46^. While maintenance of homeostasis requires multiple factors, here we focused on the neuropeptide galanin, as it has been shown to impact whole organism activity ^10–12^, sleep ^12–16^ and responses to stress ^17,20,21,47,48^. In animals such as zebrafish, sleep is operationally defined as periods of locomotor quiescence with an increased arousal threshold ^49–52^. A recent study showed evidence for galanin being important in homeostatic sleep regulation in zebrafish ^12^. Others found a decrease in locomotor activity upon galanin overexpression ^10,11^. We similarly found that overexpression of galanin leads to a reduction in brain activity, while lack of galanin leads to neuronal hyperactivity. Interestingly, most sleep-active GABAergic neurons in the ventrolateral preoptic area (VLPO) are galanin positive in mammals and the VLPO has long been proposed to be a major sleep regulator ^14,53,54^. While our investigation did not delve into galanin’s direct influence on sleep, our results suggests that galanin exerts a sedative influence on the brain. This calming effect is likely governed by *galr1a*, as lack of *galr1a* leads to similar neuronal hyperactivity of the brain as the absence of galanin. This is in line with the observation that GALR1 activation inhibits the expressing cell by opening of G protein-regulated inwardly rectifying K^+^ channels (GIRKs) ^1^. Moreover, it has been shown that galanin signaling is implicated in the reduction of glutamate release ^55–57^, which could further explain its calming effect on the brain.

Besides galanin regulating global neuronal activity, we showed that excessive neuronal activity is a potential predictor of subsequent reduced neuronal inactivity similar to another study ^12^. Employing two distinct models characterized by increased neuronal activity and epileptic seizures, our findings elucidate a consistent pattern – elevated galanin levels and decline of overall neuronal activity. This discovery is particularly interesting, considering that post-ictal fatigue is a common symptom in epilepsy patients ^58,59^ and it further mimics slow background brain activity and reduced muscle tone present in human patients ^60–62^. Moreover, the reduced neuronal activity is particularly intriguing, especially when considering the divergent mechanisms of action utilized by the two models we employed. In *eaat2a*^*-/-*^ mutants glutamate uptake into astroglia is deprecated. This results in an accumulation of glutamate in the synaptic cleft, leading to the initiation of severe epileptic seizures that spread across the entire brain 36. PTZ, on the other hand, acts as a GABAA receptor antagonist, leading to the disinhibition of the brain and a subsequent dose-dependent increase in neuronal activity, ultimately triggering epileptic seizures. The observation that both models show a reduced neuronal activity interictally (*eaat2a*^*-/-*^) or after washout (PTZ), which coincides with upregulation of galanin, suggests possible implications of galanin in various types of epileptic seizures. Galanin has been shown to be antiepileptic in numerous studies ^31–34,55^. It should also be noted that Galanin is known to be depleted by seizures in the hippocampus of mice^63^. In an attempt to assess potential antiepileptic effects of galanin in zebrafish, we manipulated galanin levels in *eaat2a*^*-/-*^ mutants. However, the impact of galanin on seizures in *eaat2a*^*-/-*^ larvae was found to be minimal, likely attributed to the exceptionally severe seizure phenotype associated with *eaat2a* dysfunction ^36^. In acute PTZ exposure however, galanin modulated seizures significantly. While we hypothesized to see antiepileptic effects upon modulation of galanin, our observations did not initially support this expectation. Galanin overexpression significantly elevated the number of seizures, whereas the absence of galanin led to a decrease in seizure occurrence. This is however in line with results from a recent publication, where galanin was found to be involved in fine-tuning a stress response upon exposure to hypertonic solution ^17^. Supposedly, galanin binds to autoreceptors on Gal^+^ neurons, reducing their own activity and therefore reducing their release of GABA on downstream stress-promoting neurons, ultimately leading to an increase in stress via Crh^+^ neurons ^17^. This can be thought of as a control system to fine-tune neuroendocrine and behavioral responses to stressful situations. While it is not clear which galanin receptor is governing this inhibitory loop, *galr1a* stands out as the most expressed and main inhibitory galanin receptor in the zebrafish brain ^41–43^. Hence, we explored whether *galr1a* crispants exhibit a reduction in seizure occurrence similar to *gal*^*-/-*^ mutants upon PTZ-induced stress. Surprisingly, we found that seizures did not decrease in number in *galr1a* crispants, indicating that *galr1a* might not function as the autoreceptor on GABAergic Gal^+^ neurons. Furthermore, seizures got more severe in *galr1a* crispants, as their amplitude and duration increased significantly, similar as in *gal*^*-/-*^ mutants. This suggests that galanin influences whole brain activity in at least two ways during epileptic seizures. On one hand, galanin acts as a sedating agent, decreasing seizure severity, which is likely governed by *galr1a*. On the other hand, galanin increases seizure occurrence, similar to increasing stress in another study ^17^ and we hypothesize that this might be dependent on one of the other galanin receptors.

Taken together, our data sheds light on a multifaceted role of galanin in regulating whole brain activity. The complexity of galanin’s effects is evident in our study, where it can serve as a central nervous system depressant or regulate stress response pathways. This is particularly the case during acute stress, where its modulatory role overrides its sedative actions. This suggests a nuanced interplay between galanin and various physiological processes, elucidating on its potential significance in modulating stress-related pathways. Exploring the mechanisms underlying these effects could offer valuable insights into the intricate role of galanin and its implications for various physiological functions or neurological disorders such as epilepsy. Other studies have revealed that galanin-producing neurons in the hypothalamus are involved in regulating diverse processes including food intake ^22–26^, parental behavior ^27,28^, sleep ^12–16^, and stress ^17–21^. Moreover, discrete subsets of hypothalamic neurons express galanin, with each subset displaying distinct responses to various stimuli. This diversity in neuronal responsiveness may account for the broad spectrum of physiological functions governed by galanin within the brain ^17^. Investigating the function of these galaninergic neuron populations represents a crucial next step in unraveling how galanin modulates neuronal activity and ultimately influences brain function. Although many questions remain, our data reveals a multifaceted role of galanin, where galanin regulates whole-brain activity but also shapes acute responses to stress and epileptic seizures.

## Materials & Methods

### Zebrafish lines and maintenance

All zebrafish experiments were performed on larvae at 5 day post fertilization (dpf). Zebrafish were kept under standard conditions at 28°C on a 14 hr/10 hr light/dark cycle ^64^. For experiments, adult animals were set up pairwise and embryos and larvae were raised in E3 medium (5 mM NaCl, 0.17 mM KCl, 0.33 mM CaCl2, 0.33 mM MgSO4). All experiments were conducted in accordance with local authorities (Zürich, Kantonales Veterinäramt TV4206). The study’s developmental stages preclude the determination of zebrafish sex. Animals were randomly allocated to experimental groups.

The following previously established transgenic lines were used in this study and combined according to needs:

*Tg(elavl3:GCaMP5G)* ^*65*^; *Tg(elavl3:GCaMP6f)* ^*66*^; *Tg(hsp70l:gal)*^*a135* 10^; *eaat2a*^*zh7 36*^ ; *gal*^*t12ae 41*^

Unless otherwise stated, experiments were performed with *nacre*^*-/-*^ (Ca^2+^ imaging and qPCR) and AB strain (qPCR) zebrafish.

### Genotyping

Larvae were genotyped after each experiment for *hsp70l:gal, eaat2a*, and *galr1a*. For experiments with *gal*^*t12ae/t12ae*^ mutants, adults were genotyped, kept in homozygosity and subsequently incrossed for experiments. For genotyping, whole larvae were anesthetized on ice, lysed in base solution (25 mM NaOH, 0.2 mM EDTA) and neutralized in neutralization solution (40 mM Tris-HCl), before PCR was performed. The target site for *hsp70l:gal* (primers: 5’-CTTGTTGACTAGAAAAATCCTTTCA-3’ and 5’-TCTCTCTTTCCTGCCAGTCC-3’), *eaat2a (*primers: 5’-GATGCAGTCGTATGGGAA-3’ and 5’-CCTTCTCCCAGATTCTCC-3’), and *galr1a (*primers: 5’-ATGGGCACCCAAAACAACAGT-3’ and 5’-GCGGCAGCACATATCCAAAAA-3’) was PCR amplified with a fast-cycling polymerase (KAPA2G Fast HotStart PCR kit, KAPA Biosystems) and a subsequent gel-electrophoresis, to allow detection of the mutant fragments.

### Quantitative Real-Time PCR (qPCR)

5 dpf larvae were anesthetized on ice and brains dissected using an insect pin and a syringe needle in a dish containing RNA later (Sigma-Aldrich). To confirm genotype, remaining tissue was lysed and gDNA amplified by means of PCR (KAPA Biosystems) as described above. Total RNA of equal pools of (n = 15) larval brains was extracted using the RNeasy kit (Qiagen). RNA was reverse transcribed to cDNA using the Super Script III First-strand synthesis system (Invitrogen). The qPCR reactions were performed using SsoAdvanced Universal SYBR Green Supermix on a CFX96 Touch Real-Time PCR Detection System (Bio-Rad). *actb1, tbp* and *rpl13a* were chosen as reference genes. Primer efficiencies for *gal* (5’-TGAGGATGTCGTCCATACCATC-3’ and 5’-GGTTGACTGATCTCTTCTGATGTG-3’), *actb1* (5’-CAGACATCAGGGAGTGATGGTTGG-3’ and 5’-CAGATCTTCTCCATGTCATCCCAG-3’), *tbp* (5’-ACAACAGCCTACCTCCTTTCG-3’ and 5’-CGTCCCATACGGCATCATAGG-3’) and *rpl13a* (5’-TCTGGAGGACTGTAAGAGGTATGC-3’ and 5’-AGACGCACAATCTTGAGAGCAG-3’ ^67^) were calculated by carrying out a dilution series and specificity was determined using melting curve analysis. After primer efficiencies were determined equal, brain samples were used for qPCR using 1 ng of cDNA per reaction. The controls “no reverse transcription control” (nRT control) and “no template control” (NTC) were performed with every qPCR reaction. All reactions were performed in technical triplicates. Data was analyzed in CFX Maestro Software from Bio-Rad and GraphPad Prism version 10.1.2 for Windows, GraphPad Software, Boston, Massachusetts USA, www.graphpad.com. Expression of *gal* was normalized to expression of *actb1, tbp* and *rpl13a*. Relative expression levels were calculated using the ΔΔCt method. Each experiment was performed in 3 biological replicates.

### F0 Knockouts

We performed F0 knockouts similar as described in Kroll et al ^44,68,69^. In brief, 3 CRISPR target sites for *galr1a* (*ENSDARG00000005522*) were selected with CHOPCHOP ^70^. While it was not possible to generate efficient guides targeting 3 different exons, we chose to generate 3 guide RNAs (#1 (5’-CGCCTACTATCAGGGCATCGTGG-3’), #2 (5’-TGGCGTCCTTGGAAACTCTCTGG-3’), #3 (5’-GGTGAAGATGCTTACGAGCATGG-3’) targeting the same exon (exon 1). Reagents were ordered from integrated DNA technologies (IDT) and were resuspended, annealed and pooled according to Kroll et al ^68^. For generating the F0 knockouts, one-cell staged embryos were injected with 1 nl injection mix (10.1 fmol [357 pg] per gRNA) into the yolk. Control injection mix contained a set of three scrambled RNPs, which carry gRNAs (IDT, Alt-R CRISPR-Cas9 Negative Control crRNA #1 (#1072544), #2 (#1072545), #3 (#1072546)^44^) with pseudo-random sequences predicted to not match any genomic locus. For all experiments scramble RNP injected siblings were used as controls. As stable *galr1a* mutants ^41^ lack overt morphological phenotypes apart from their pigmentation phenotype, all larvae with abnormal appearance were discarded to rule out injection artifacts.

### Calcium imaging and data analysis

5 dpf larvae in the Tg(*elavl3:GCaMP5G*) or Tg(*elavl3:GCaMP6f*) background respectively were individually immobilized in a drop of 2% low melting temperature agarose (Sigma-Aldrich) in a small cell culture dish. Agarose was left to dry briefly before a part was cut away and a second larvae was embedded close to the first larvae and in the same focal plane. Dish was filled with E3 medium. Calcium signals were recorded using an Olympus BX51 WI epifluorescence microscope with a 5x objective and by means of a camera (1.33 Hz sampling rate) for 1 h with the VisiView software established by the Visitron Systems GmbH. In each experiment, the GCaMP fluorescence signal of manually selected regions of interest (ROIs, whole brain) was extracted using Fiji ImageJ ^71^. Images were corrected for sample drift with Correct_3D_Drift.py script ^72^ in Fiji ImageJ ^71^. For each time point, the mean intensity of each ROI was measured and further processed using a custom script in R with the RStudio interface ^73,74^. The baseline (F_0_) as 1% of the total fluorescence of every ROI was computed within a moving window of 150s (for normal brain activity) or 300s (for *eaat2a*^-/-^ seizure analysis) per fish. Subsequently, the fractional change in fluorescence (ΔF/F_0_) of each ROI was determined. In all imaging experiments, heartbeat of larvae was assessed before and after each experiment, and animals without heartbeat were excluded from analysis. In order to compare whole brain activity, thresholds for event detection were set. For examining normal brain activity, events were categorized into two distinct groups, 5% and 10% of ΔF/F_0_. Duration, amplitude and number of events were calculated for each of these groups. For analyzing normal behavior, area under the curve (AUC) was calculated for the whole normalized fluorescence trace except for the experiment with *eaat2a* mutants and the respective controls in Figure 1B, where AUC was calculated and averaged over 2 five-minutes time windows per animal (same random windows for all larvae, adjusted if during seizure). To demonstrate previously published *eaat2a* hypoactivity ^36^ (Figure 1A, B, D, E, F), data from *eaat2a*^*-/-*^ (Figure 3A – F) and closely related wild type larvae (Figure 2A – F) were re-used and differently analyzed.

For this particular experiment, we removed seizures by excluding all events with amplitudes bigger than 50% ΔF/F_0_ and a duration of more than 20s. For all other recordings, seizures were defined as calcium fluctuations reaching at least 100% of ΔF/F_0_ in the whole brain. Seizure duration was defined as the period from the first time point where ΔF/F_0_ is greater than 50% (seizure initiation) until the first time point where ΔF/F_0_ is below 50% in the brain. To generate the overlay plots (Figure 3A, 3G, 4A, 4G and 4M); all seizures were aligned by the time point where 50% of the maximum ΔF/F_0_ was reached. The resulting plots showcase the mean average seizure per genotype, shaded area represents the standard error of the mean (SEM). AUC and time to peak of each seizure were calculated from the beginning of the aligned seizure trace.

### PTZ exposure

For exposure with pentylenetetrazole (PTZ), 20 mM PTZ was prepared freshly on each experimental day (Sigma-Aldrich). For acute PTZ exposure, E3 medium was replaced by 20 mM PTZ (in E3) directly before starting the 1 h imaging period. For PTZ rebound experiments, freely swimming larvae were exposed to 20 mM PTZ (in E3) in a 50 ml cell culture dish for 1 h, subsequently rinsed in a dish containing E3 medium and were then transferred to a fresh dish with E3 medium for 1 h. Larvae were embedded as described earlier and left in E3 medium until the 1-hour calcium imaging experiment was conducted precisely 2 hours after initiating the PTZ washout.

### Quantification and statistical analysis

Statistical analysis was performed using R software version 4.2.1 with the RStudio version RStudio 2023.09.1+494 interface ^73,74^ or GraphPad Prism version 10.1.2 for Windows, GraphPad Software, Boston, Massachusetts USA, www.graphpad.com. Calcium imaging data was tested for normality using Shapiro–Wilk test. Wilcoxon-Mann-Whitney test nonpaired analysis was used for calcium imaging analysis (amplitudes, duration, AUC, time to peak). For comparing event number, as data was overdispersed, a negative binomial generalized linear model was fitted and assessed (package: MASS, glm.nb in R). For qPCR data, Student’s t-test in GraphPad Prism was used. Differences were considered statistically significant if p < 0.05. Description of the number of animals used for each experiment can be found in the figures and figure legends.

### Whole-mount Immunostaining

5 dpf larvae were fixed in 4% PFA, washed in PBS and permeabilized using acetone at - 20°C. Larvae were blocked using PBDT (PBS + 1% BSA + 0.5% Triton X-100 + 1% DMSO) with 10% goat serum for 30 minutes at room temperature and subsequently incubated with primary antibodies overnight at 4°C. Rabbit anti-galanin (1:400, EDM Milipore AB5909), Mouse anti-acetylated tubulin (IgG2b, 1:500, Sigma 7451) were used as primary antibodies. Following PBDT washes, embryos were incubated in secondary antibodies (Goat anti-Rabbit Alexa Fluor 488 (Invitrogen A-11008, ThermoFisher Scientific) and goat anti-mouse IgG2b Alexa Fluor 647 (Invitrogen A-21242, ThermoFisher Scientific)) for at least 2 hours at room temperature. Larvae were cleared in an increasing glycerol row and stored in 70% glycerol in PBS. Larval heads were cut and mounted using Mowiol (Polysciences).

Confocal images were acquired using the Leica TCS LSI microscope. Maximum z-projections were created using Fiji ImageJ ^71^.

## Acknowledgments

We thank P. Podlasz (University of Warmia and Mazury, Poland) and U. Irion (Max Planck Institute, Tübingen) for transgenic lines. We thank M. Walther, K. Dannenhauer and H. Möckel for excellent technical and animal support and the Neuhauss and Bachmann lab members for valuable discussions. This work was funded by the Swiss National Science Foundation (SNF 31003A_135598, 310030_204648) to NNR, MR, ALH and SCFN and UZH Forschungskredit Candoc Grant K-74417-04-01 to NNR.

---

**Supplementary Figure 1.**
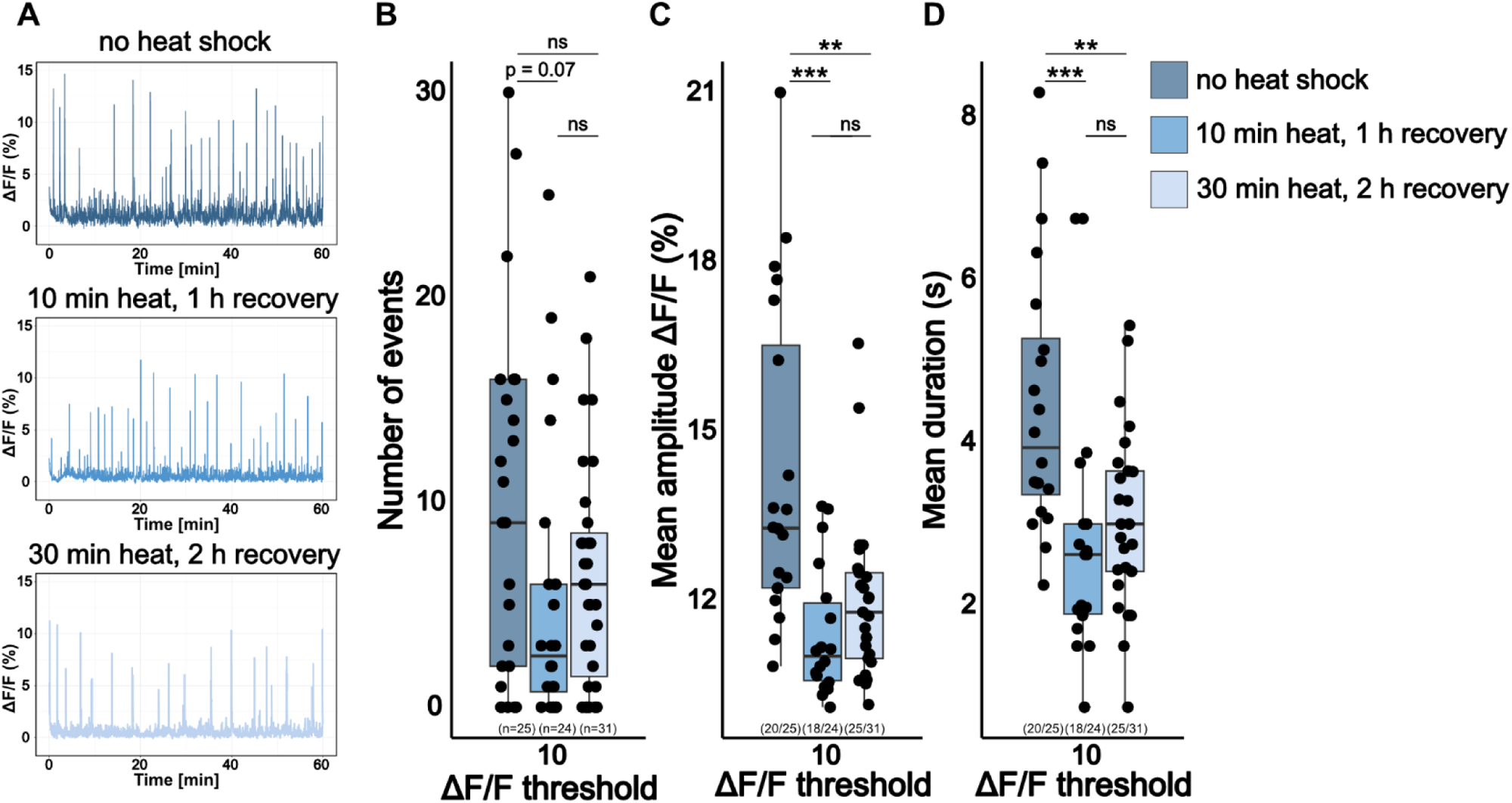
Heat Shock Decreases Brain Activity of 5 dpf Wild Type Larvae. A) Representative calcium signals (*elavl3:GCaMP5G*) recorded across the brain of 5 dpf unexposed wild type larva (blue, top) and 5 dpf wild type larvae with a 37°C heat shock for either 10 min (steelblue blue, middle) or 30 min (light blue, bottom) with different recovery times at room temperature. Significance levels: ***p < .001, **p < .01, *p < .05, ns = not significant (p > .05), Wilcoxon-Mann-Whitney test (C, D), negative binomial regression (B).

**Supplementary Figure 2.**
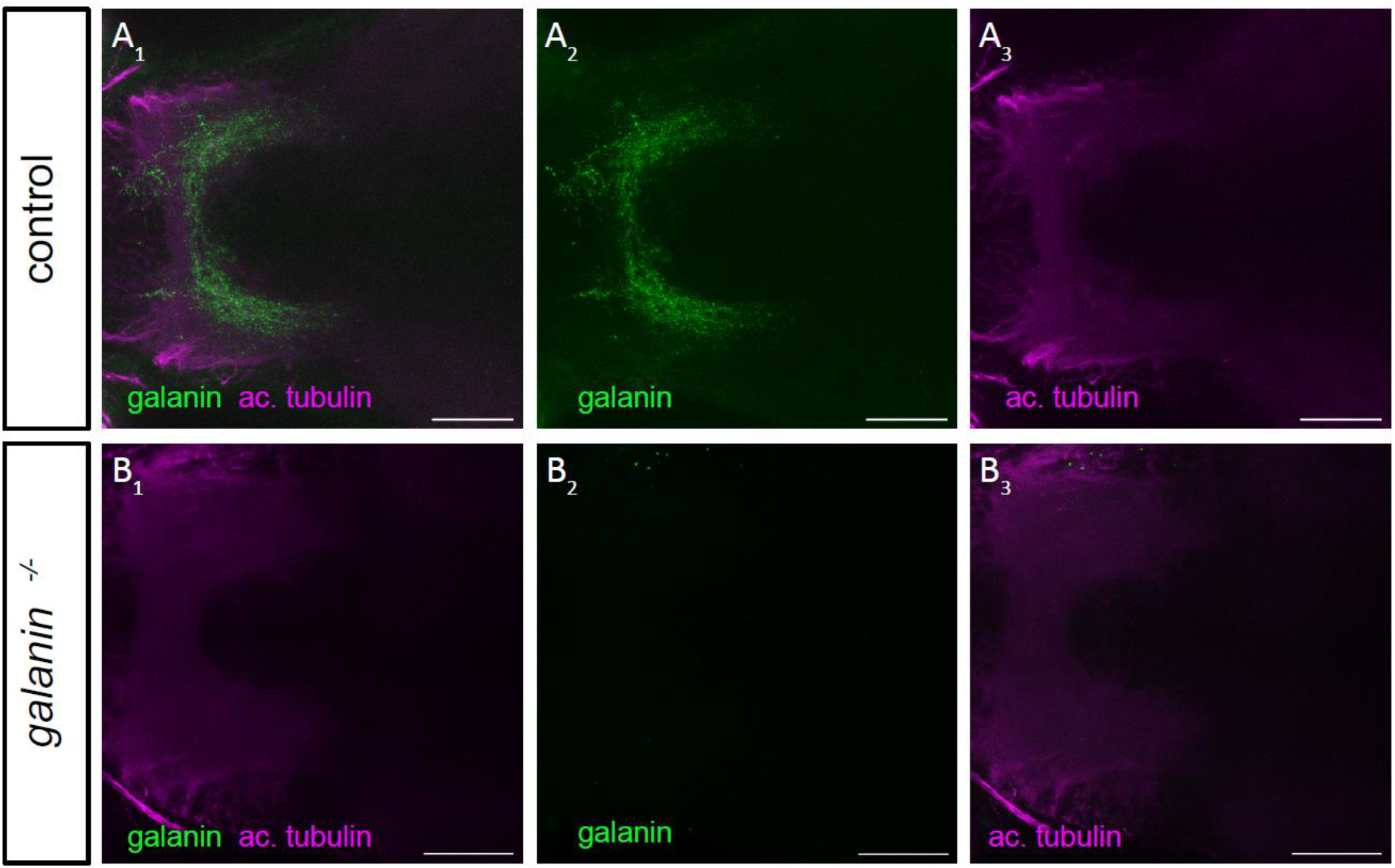
Immunostaining Confirms Protein Deficiency in *gal* ^*-/-*^ Mutants. Whole-mount Immunostaining of galanin localized in the diencephalon of 5 dpf zebrafish larvae. Antibodies used for staining included anti-galanin (green) and anti-ac. tubulin (acetylated tubulin, magenta). A1) representative galanin staining in control larvae. A2,A3) Single immunohistochemical channels. B1) representative galanin staining in *gal* ^*-/-*^ larvae. B2, B3) Single immunohistochemical channels. Scale bar = 25 μm

**Supplementary movie 1 - Whole brain activity recording in control and *hsp70:gal* larvae**

Representative raw data of recorded calcium signals (*elavl3:GCaMP5G*) recorded across the brain of 5 dpf control (left) and *hsp70:gal* (right, galanin-OE) over a 15 min time period chosen at random. The recordings were exported with 7fps. Both larvae were recorded on the same day.

**Supplementary movie 2 - Whole brain activity recording in control and *gal***^***-/-***^ **larvae**

Representative raw data of recorded calcium signals (*elavl3:GCaMP5G*) recorded across the brain of 5 dpf control (left) and *gal*^*-/-*^ (right, galanin-KO) over a 15 min time period chosen at random. The recordings were exported with 7fps. Both larvae were recorded on the same day.

**Supplementary movie 3 - Whole brain activity recording in control and galr1aKO larvae**

Representative raw data of recorded calcium signals (*elavl3:GCaMP5G*) recorded across the brain of 5 dpf control (left) and galr1aKO (right) over a 15 min time period chosen at random. The recordings were exported with 7fps. Both larvae were recorded on the same day.

**Supplementary movie 4 – Seizure recordings in control and *hsp70:gal* larvae in *eaat2a*** ^***-/-***^ **background**

Representative raw data of recorded calcium signals (*elavl3:GCaMP5G*) during a typical seizure recorded across the brain of 5 dpf control (left) and *hsp70:gal* (right, galanin-OE) in *eaat2a* ^*-/-*^ background. The seizures were captured over 150 frames (about 1 minute 50 seconds recording) and exported with 7fps. Both larvae were recorded on the same day.

**Supplementary movie 5 – Seizure recordings in control and *gal***^***-/-***^ **larvae in *eaat2a*** ^***-/-***^ **background**

Representative raw data of recorded calcium signals (*elavl3:GCaMP5G*) during a typical seizure recorded across the brain of 5 dpf control (left) and *gal*^*-/-*^ (right, galanin-KO) in *eaat2a* ^*-/-*^ background. The seizures were captured over 150 frames (about 1 minute 50 seconds recording) and exported with 7fps. Both larvae were recorded on the same day.

**Supplementary movie 6 – Seizure recordings in control and *hsp70:gal* larvae acutely exposed to 20mM PTZ**

Representative raw data of recorded calcium signals (*elavl3:GCaMP5G*) during a typical seizure recorded across the brain of 5 dpf control (left) and *hsp70:gal* (right, galanin-OE) acutely exposed to 20mM PTZ. The seizures were captured over 125 frames (about 1 minute 50 seconds recording) and exported with 7fps. Both larvae were recorded on the same day.

**Supplementary movie 7 – Seizure recordings in control and *gal***^***-/-***^ **larvae acutely exposed to 20mM PTZ**

Representative raw data of recorded calcium signals (*elavl3:GCaMP5G*) during a typical seizure recorded across the brain of 5 dpf control (left) and *gal*^*-/-*^ (right, galanin-KO) acutely exposed to 20mM PTZ. The seizures were captured over 125 frames (about 1 minute 50 seconds recording) and exported with 7fps. Both larvae were recorded on the same day.

**Supplementary movie 8 – Seizure recordings in control and galr1aKO larvae acutely exposed to 20mM PTZ**

Representative raw data of recorded calcium signals (*elavl3:GCaMP5G*) during a typical seizure recorded across the brain of 5 dpf control (left) and galr1aKO (right) acutely exposed to 20mM PTZ. The seizures were captured over 125 frames (about 1 minute 50 seconds recording) and exported with 7fps. Both larvae were recorded on the same day.

## References

1. Lang, R. et al. Physiology, signaling, and pharmacology of galanin peptides and receptors: three decades of emerging diversity. Pharmacological reviews 67, 118–175; 10.1124/pr.112.006536 (2015).

2. van den Pol, Anthony N. Neuropeptide Transmission in Brain Circuits. Neuron 76, 98–115; 10.1016/j.neuron.2012.09.014 (2012).

3. Purves, D. et al. (eds.). Neuroscience (Sinauer Associates Oxford University Press, New York, Oxford, 2019).

4. Douglas, W. W. Stimulus-secretion coupling: the concept and clues from chromaffin and other cells. British journal of pharmacology 34, 451–474; 10.1111/j.1476-5381.1968.tb08474.x (1968).

5. MacArthur, L. & Eiden, L. Neuropeptide genes: Targets of activity-dependent signal transduction. Peptides 17, 721–728; 10.1016/0196-9781(95)02100-0 (1996).

6. Jan, L. Y. & Jan, Y. N. Peptidergic transmission in sympathetic ganglia of the frog. The Journal of physiology 327, 219–246; 10.1113/jphysiol.1982.sp014228 (1982).

7. Atkinson, L. E. et al. Ascaris suum Informs Extrasynaptic Volume Transmission in Nematodes. ACS chemical neuroscience 12, 3176–3188; 10.1021/acschemneuro.1c00281 (2021).

8. Nässel, D. R. Neuropeptide signaling near and far: how localized and timed is the action of neuropeptides in brain circuits? Invertebrate neuroscience : IN 9, 57–75; 10.1007/s10158-009-0090-1 (2009).

9. Ripoll-Sánchez, L. et al. The neuropeptidergic connectome of C. elegans. Neuron 111, 3570-3589.e5; 10.1016/j.neuron.2023.09.043 (2023).

10. Woods, I. G. et al. Neuropeptidergic signaling partitions arousal behaviors in zebrafish. The Journal of neuroscience : the official journal of the Society for Neuroscience 34, 3142–3160; 10.1523/JNEUROSCI.3529-13.2014 (2014).

11. Podlasz, P., Jakimiuk, A., Kasica-Jarosz, N., Czaja, K. & Wasowicz, K. Neuroanatomical Localization of Galanin in Zebrafish Telencephalon and Anticonvulsant Effect of Galanin Overexpression. ACS chemical neuroscience 9, 3049–3059; 10.1021/acschemneuro.8b00239 (2018).

12. Reichert, S., Pavón Arocas, O. & Rihel, J. The Neuropeptide Galanin Is Required for Homeostatic Rebound Sleep following Increased Neuronal Activity. Neuron 104, 370-384.e5; 10.1016/j.neuron.2019.08.010 (2019).

13. Gaus, S. E., Strecker, R. E., Tate, B. A., Parker, R. A. & Saper, C. B. Ventrolateral preoptic nucleus contains sleep-active, galaninergic neurons in multiple mammalian species. Neuroscience 115, 285–294; 10.1016/s0306-4522(02)00308-1 (2002).

14. Kroeger, D. et al. Galanin neurons in the ventrolateral preoptic area promote sleep and heat loss in mice. Nature communications 9, 4129; 10.1038/s41467-018-06590-7 (2018).

15. McGinty, D. & Szymusiak, R. Hypothalamic regulation of sleep and arousal. Frontiers in bioscience : a journal and virtual library 8; 10.2741/1159 (2003).

16. Ma, Y. et al. Galanin Neurons Unite Sleep Homeostasis and α2-Adrenergic Sedation. Current Biology 29, 3315-3322.e3; 10.1016/j.cub.2019.07.087 (2019).

17. Corradi, L., Bruzzone, M., Maschio, M. d., Sawamiphak, S. & Filosa, A. Hypothalamic Galanin-producing neurons regulate stress in zebrafish through a peptidergic, self-inhibitory loop. Current Biology 32, 1497-1510.e5; 10.1016/j.cub.2022.02.011 (2022).

18. Juhasz, G. et al. Brain galanin system genes interact with life stresses in depression-related phenotypes. Proceedings of the National Academy of Sciences of the United States of America 111, E1666–73; 10.1073/pnas.1403649111 (2014).

19. Hökfelt, T. et al. Neuropeptide and Small Transmitter Coexistence: Fundamental Studies and Relevance to Mental Illness. Frontiers in neural circuits 12, 106; 10.3389/fncir.2018.00106 (2018).

20. Khoshbouei, H., Cecchi, M. & Morilak, D. Modulatory effects of galanin in the lateral bed nucleus of the stria terminalis on behavioral and neuroendocrine responses to acute stress. Neuropsychopharmacology : official publication of the American College of Neuropsychopharmacology 27, 25–34; 10.1016/S0893-133X(01)00424-9 (2002).

21. Picciotto, M. R., Brabant, C., Einstein, E. B., Kamens, H. M. & Neugebauer, N. M. Effects of galanin on monoaminergic systems and HPA axis: Potential mechanisms underlying the effects of galanin on addiction- and stress-related behaviors. Brain Research 1314, 206–218; 10.1016/j.brainres.2009.08.033 (2010).

22. Schick, R. R. et al. Effect of galanin on food intake in rats: involvement of lateral and ventromedial hypothalamic sites. American Journal of Physiology-Regulatory, Integrative and Comparative Physiology; 10.1152/ajpregu.1993.264.2.R355.

23. Adams, A. C., Clapham, J. C., Wynick, D. & Speakman, J. Feeding Behaviour in Galanin Knockout Mice Supports a Role of Galanin in Fat Intake and Preference. Journal of Neuroendocrinology 20, 199–206; 10.1111/j.1365-2826.2007.01638.x (2008).

24. Qualls-Creekmore, E. et al. Galanin-Expressing GABA Neurons in the Lateral Hypothalamus Modulate Food Reward and Noncompulsive Locomotion. J. Neurosci. 37, 6053–6065; 10.1523/JNEUROSCI.0155-17.2017 (2017).

25. Laque, A. et al. Leptin modulates nutrient reward via inhibitory galanin action on orexin neurons. Molecular Metabolism 4, 706–717; 10.1016/j.molmet.2015.07.002 (2015).

26. Leibowitz, S. F., Akabayashi, A. & Wang, J. Obesity on a High-Fat Diet: Role of Hypothalamic Galanin in Neurons of the Anterior Paraventricular Nucleus Projecting to the Median Eminence. J. Neurosci. 18, 2709–2719; 10.1523/JNEUROSCI.18-07-02709.1998 (1998).

27. Kohl, J. et al. Functional circuit architecture underlying parental behaviour. Nature 556, 326–331; 10.1038/s41586-018-0027-0 (2018).

28. Wu, Z., Autry, A. E., Bergan, J. F., Watabe-Uchida, M. & Dulac, C. G. Galanin neurons in the medial preoptic area govern parental behaviour. Nature 509, 325–330; 10.1038/nature13307 (2014).

29. Kovac, S. & Walker, M. C. Neuropeptides in epilepsy. Neuropeptides 47, 467–475; 10.1016/j.npep.2013.10.015 (2013).

30. Lerner, J. T., Sankar, R. & Mazarati, A. M. Galanin and epilepsy. Cellular and molecular life sciences : CMLS 65, 1864–1871; 10.1007/s00018-008-8161-8 (2008).

31. McColl, C. D., Jacoby, A. S., Shine, J., Iismaa, T. P. & Bekkers, J. M. Galanin receptor-1 knockout mice exhibit spontaneous epilepsy, abnormal EEGs and altered inhibition in the hippocampus. Neuropharmacology 50, 209–218; 10.1016/j.neuropharm.2005.09.001 (2006).

32. Fetissov, S. O. et al. Altered hippocampal expression of neuropeptides in seizure-prone GALR1 knockout mice. Epilepsia 44, 1022–1033; 10.1046/j.1528-1157.2003.51402.x (2003).

33. Jacoby, A. S., Hort, Y. J., Constantinescu, G., Shine, J. & Iismaa, T. P. Critical role for GALR1 galanin receptor in galanin regulation of neuroendocrine function and seizure activity. Brain research. Molecular brain research 107; 10.1016/s0169-328x(02)00451-5 (2002).

34. Mazarati, A. M. et al. Modulation of Hippocampal Excitability and Seizures by Galanin. J. Neurosci. 20, 6276–6281; 10.1523/JNEUROSCI.20-16-06276.2000 (2000).

35. Drexel, M., Locker, F., Kofler, B. & Sperk, G. Effects of galanin receptor 2 and receptor 3 knockout in mouse models of acute seizures. Epilepsia 59, e166–e171; 10.1111/epi.14573 (2018).

36. Hotz, A. L. et al. Loss of glutamate transporter eaat2a leads to aberrant neuronal excitability, recurrent epileptic seizures, and basal hypoactivity. Glia 70, 196–214; 10.1002/glia.24106 (2022).

37. Baraban, S. C., Dinday, M. T. & Hortopan, G. A. Drug screening in Scn1a zebrafish mutant identifies clemizole as a potential Dravet syndrome treatment. Nature communications 4, 2410; 10.1038/ncomms3410 (2013).

38. Hortopan, G. A., Dinday, M. T. & Baraban, S. C. Spontaneous seizures and altered gene expression in GABA signaling pathways in a mind bomb mutant zebrafish. The Journal of neuroscience : the official journal of the Society for Neuroscience 30, 13718–13728; 10.1523/JNEUROSCI.1887-10.2010 (2010).

39. Baraban, S. C., Taylor, M. R., Castro, P. & Baier, H. Pentylenetetrazole induced changes in zebrafish behavior, neural activity and c-fos expression. Neuroscience 131, 759–768; 10.1016/j.neuroscience.2004.11.031 (2005).

40. Baraban, S. C. et al. A large-scale mutagenesis screen to identify seizure-resistant zebrafish. Epilepsia 48, 1151–1157; 10.1111/j.1528-1167.2007.01075.x (2007).

41. Eskova, A., Frohnhöfer, H. G., Nüsslein-Volhard, C. & Irion, U. Galanin Signaling in the Brain Regulates Color Pattern Formation in Zebrafish. Current biology : CB 30, 298-303.e3; 10.1016/j.cub.2019.11.033 (2020).

42. Li, L. et al. A novel galanin receptor 1a gene in zebrafish: tissue distribution, developmental expression roles in nutrition regulation. Comparative biochemistry and physiology. Part B, Biochemistry & molecular biology 164, 159–167; 10.1016/j.cbpb.2012.12.004 (2013).

43. Kim, D.-K. et al. Coevolution of the spexin/galanin/kisspeptin family: Spexin activates galanin receptor type II and III. Endocrinology 155, 1864–1873; 10.1210/en.2013-2106 (2014).

44. Kroll, F. et al. A simple and effective F0 knockout method for rapid screening of behaviour and other complex phenotypes. eLife 10; 10.7554/eLife.59683 (2021).

45. Ma, X. et al. Effects of galanin receptor agonists on locus coeruleus neurons. Brain Research 919, 169–174; 10.1016/S0006-8993(01)03033-5 (2001).

46. Lin, A. et al. Imaging whole-brain activity to understand behavior. Nature reviews. Physics 4, 292–305; 10.1038/s42254-022-00430-w (2022).

47. Tillage, R. P. et al. Co-released norepinephrine and galanin act on different timescales to promote stress-induced anxiety-like behavior. Neuropsychopharmacol. 46, 1535–1543; 10.1038/s41386-021-01011-8 (2021).

48. Tillage, R. P., Wilson, G. E., Liles, L. C., Holmes, P. V. & Weinshenker, D. Chronic Environmental or Genetic Elevation of Galanin in Noradrenergic Neurons Confers Stress Resilience in Mice. The Journal of neuroscience : the official journal of the Society for Neuroscience 40, 7464–7474; 10.1523/JNEUROSCI.0973-20.2020 (2020).

49. Barlow, I. L. & Rihel, J. Zebrafish sleep: from geneZZZ to neuronZZZ. Current Opinion in Neurobiology 44, 65–71; 10.1016/j.conb.2017.02.009 (2017).

50. Hendricks, J. C. et al. Rest in Drosophila Is a Sleep-like State. Neuron 25, 129–138; 10.1016/s0896-6273(00)80877-6 (2000).

51. Raizen, D. M. et al. Lethargus is a Caenorhabditis elegans sleep-like state. Nature 451, 569–572; 10.1038/nature06535 (2008).

52. van Alphen, B., Yap, M. H., Kirszenblat, L., Kottler, B. & van Swinderen, B. A Dynamic Deep Sleep Stage in Drosophila. J. Neurosci. 33, 6917–6927; 10.1523/JNEUROSCI.0061-13.2013 (2013).

53. Gaus, S., Strecker, R., Tate, B., Parker, R. & Saper, C. Ventrolateral preoptic nucleus contains sleep-active, galaninergic neurons in multiple mammalian species. Neuroscience 115, 285–294; 10.1016/s0306-4522(02)00308-1 (2002).

54. Lu, J., Greco, M. A., Shiromani, P. & Saper, C. B. Effect of Lesions of the Ventrolateral Preoptic Nucleus on NREM and REM Sleep. J. Neurosci. 20, 3830–3842; 10.1523/JNEUROSCI.20-10-03830.2000 (2000).

55. Kokaia, M. et al. Suppressed kindling epileptogenesis in mice with ectopic overexpression of galanin. Proceedings of the National Academy of Sciences of the United States of America 98, 14006–14011; 10.1073/pnas.231496298 (2001).

56. Elliott-Hunt, C. R. et al. Galanin acts as a neuroprotective factor to the hippocampus. Proceedings of the National Academy of Sciences of the United States of America 101, 5105–5110; 10.1073/pnas.0304823101 (2004).

57. Zini, S., Roisin, M.-P., Langel, U., Bartfai, T. & Ben-Ari, Y. Galanin reduces release of endogeneous excitatory amino acids in the rat hippocampus. European Journal of Pharmacology: Molecular Pharmacology 245, 1–7; 10.1016/0922-4106(93)90162-3 (1993).

58. Hamelin, S., Kahane, P. & Vercueil, L. Fatigue in epilepsy: a prospective interictal and post-ictal survey. Epilepsy research 91, 153–160; 10.1016/j.eplepsyres.2010.07.006 (2010).

59. Ettinger, A. B., Weisbrot, D. M., Nolan, E. & Devinsky, O. Postictal symptoms help distinguish patients with epileptic seizures from those with non-epileptic seizures. Seizure 8, 149–151; 10.1053/seiz.1999.0270 (1999).

60. Allen, A. S. et al. De novo mutations in epileptic encephalopathies. Nature 501, 217–221; 10.1038/nature12439 (2013).

61. Epi4K Consortium. De Novo Mutations in SLC1A2 and CACNA1A Are Important Causes of Epileptic Encephalopathies. American journal of human genetics 99, 287–298; 10.1016/j.ajhg.2016.06.003 (2016).

62. Guella, I. et al. De Novo Mutations in YWHAG Cause Early-Onset Epilepsy. American journal of human genetics 101, 300–310; 10.1016/j.ajhg.2017.07.004 (2017).

63. Mazarati, A. M. et al. Galanin modulation of seizures and seizure modulation of hippocampal galanin in animal models of status epilepticus. J. Neurosci. 18, 10070–10077; 10.1523/JNEUROSCI.18-23-10070.1998 (1998).

64. Mullins, M. C., Hammerschmidt, M., Haffter, P. & Nüsslein-Volhard, C. Large-scale mutagenesis in the zebrafish: in search of genes controlling development in a vertebrate. Current Biology 4, 189–202; 10.1016/s0960-9822(00)00048-8 (1994).

65. Akerboom, J. et al. Optimization of a GCaMP calcium indicator for neural activity imaging. J. Neurosci. 32, 13819–13840; 10.1523/JNEUROSCI.2601-12.2012 (2012).

66. Chen, T.-W. et al. Ultrasensitive fluorescent proteins for imaging neuronal activity. Nature 499, 295–300; 10.1038/nature12354 (2013).

67. Lin, C., Spikings, E., Zhang, T. & Rawson, D. Housekeeping genes for cryopreservation studies on zebrafish embryos and blastomeres. Theriogenology 71, 1147–1155; 10.1016/j.theriogenology.2008.12.013 (2009).

68. Kroll, F. & Rihel, J. F0 knockout—single gene v2 (2021).

69. Kroll, F. How to select the best gRNA(s) for frameshift knockouts in zebrafish v1 (2022).

70. Labun, K. et al. CHOPCHOP v3: expanding the CRISPR web toolbox beyond genome editing. Nucleic acids research 47, W171–W174; 10.1093/nar/gkz365 (2019).

71. Schindelin, J. et al. Fiji: an open-source platform for biological-image analysis. Nature methods 9, 676–682; 10.1038/nmeth.2019 (2012).

72. Parslow, A., Cardona, A. & Bryson-Richardson, R. J. Sample drift correction following 4D confocal time-lapse imaging. Journal of visualized experiments : JoVE; 10.3791/51086 (2014).

73. R Core Team. R: A language and environment for statistical computing. (2022).

74. Posit team. RStudio: Integrated Development Environment for R (2023).

